# Putting hands to rest: efficient deep CNN-RNN architecture for chemical named entity recognition with no hand-crafted rules

**DOI:** 10.1101/321224

**Authors:** Ilia Korvigo, Maxim Holmatov, Anatolii Zaikovskii, Mikhail Skoblov

## Abstract

Chemical named entity recognition (NER) is an active field of research in biomedical natural language processing. To facilitate the development of new and superior chemical NER systems, BioCreative released the CHEMDNER corpus, an extensive dataset of diverse manually annotated chemical entities. Most of the systems trained on the corpus rely on complicated hand-crafted rules or curated databases for data preprocessing, feature extraction and output post-processing, though modern machine learning algorithms, such as deep neural networks, can automatically design the rules with little to none human intervention. Here we explored this approach by experimenting with various deep learning architectures for targeted tokenisation and named entity recognition. Our final model, based on a combination of convolutional and stateful recurrent neural networks with attention-like loops and hybrid word-and character-level embeddings, reaches near human-level performance on the testing dataset with no manually asserted rules. To make our model easily accessible for standalone use and integration in third-party software, we’ve developed a Python package with a minimalistic user interface.

## Content Background

Modern data-generation capabilities have clearly surpassed our capacity to manually analyse published data, which is ever-more evident in the era of high-throughput methods. Naturally, this fuels the development of automatic natural language processing (NLP) systems capable of extracting and transforming specific information from a body of literature with human-level precision. Among all the subtasks NLP introduces, named entity recognition (NER) – aiming to identify objects of particular semantic value (e.g. chemical compounds) – is one of the most fundamental for higher level event-focused analyses. Traditionally, chemical NER systems have relied on curated dictionaries and hand-crafted rules (e.g. regular expressions for systematic IUPAC names or databases of trivial names and identifiers), which are hard to develop and maintain due to diverse morphology and rich vocabulary of biomedical literature. On the other hand, various machine learning (ML) models can automatically infer efficient rules (input transformations) from annotated corpora reducing development and maintenance costs. In ML terms named entity recognition is a supervised labelling problem.

To facilitate the development of new and superior NER systems, BioCreative announced the CHEMDNER challenge, which ended in 2015 [1]. As part of this task, a team of experts has produced an extensive manually annotated corpus covering various chemical entity types, including systematic and trivial names, abbreviations and identifiers, formulae and phrases. Due to many difficulties inherent to chemical entity detection and normalisation [1], even manual annotation yields the interannotator agreement score of 91%, which can be regarded as the theoretical limit for any automatic system trained on this corpus. Twenty six teams have submitted their NER systems for the challenge, best of which have reached the F1 score of ∼ 72 − 88% [2, 3, 4, 5, 6, 7,8,9] on two subtasks: chemical entity mention (CEM) and chemical document indexing (CDI).

The systems were quite diverse in terms of text preprocessing, which is a separate NLP problem in its own right. Obviously, it’s possible to represent any text as a raw sequence of characters (e.g. byte-like sequences or Unicode character codes), yet it is more common to break the characters into word-like structures known as tokens, which can be further normalised and/or encoded. Although tokenisation typically reduces the number of time-steps in the sequence, thus reducing the input complexity, it can introduce severe artefacts, e.g. merged/overlapping entities [5, 9]. It makes it essential to use an adequate tokeniser with rules finely adjusted for the task at hand.

While there are many token encoding strategies, they all can be divided into two major groups: morphology aware (character-level) and unaware (word-level). In the latter case, one usually builds a vocabulary of all tokens occurring in a corpus and applies a minimal frequency cutoff to remove noisy entries (e.g. misspelled words and typos). Consequently, all tokens in the vocabulary get a unique identifier *t*∊ℕ_+_, while all out-of-vocabulary (OOV) tokens get a special shared identifier. The vocabulary itself can be represented as a matrix **T** = (t_1_,…, t_*T*_) of orthogonal unit vectors (also known as one-hot encodings), both sparse and purely categorical: their pair-wise distances carry no underlying information about semantical similarity. In their chemical NER system, Lu et al. [9] successfully used the skip-gram embedding model to overcome these limitations. The model uses context information and a shallow neural network to embed high-dimensional one-hot encoded vectors in a lower-dimensional vector space, wherein pair-wise distances represent semantical similarity [10, 11]. Despite this strategy’s increasing popularity, few CHEMDNER task participants have employed it for morphology unaware encoding, relying instead on manually selected features to expand token identifiers into feature vectors. While word-level encodings are efficient for morphologically rigid corpora (e.g. standard English texts), morphologically rich biomedical and chemical literature introduces many infrequent words and word-forms, resulting in high out-of-vocabulary (OOV) rates [12, 13]. Consequently, most CHEMNDER participants have additionally (or exclusively) used morphology aware-encodings, targeting various manually designed character-level features. Machine-learning models were far less diverse: since textual data are sequential, that is a value *t_i_* at time-step *i* can be conditioned on the values occurring before and after the time-step, it is only natural to use sequential models for NER problems. Although many such models exist, most of the top-scoring ML-based tools submitted for the CHEMDNER task utilised conditional random fields (CRF), which are traditionally used for sequence labelling. CRFs are graphical models related to hidden Markov models (HMMs). They take a sequence of feature vectors as inputs and generate a sequence of labels, which can be further modified during post-processing. The participants used hand-crafted post-processing rules as diverse as the preprocessing procedures.

From this brief overview of the NER systems submitted for the CHEMDNER task, it becomes quite evident that, despite the introduction of machine learning methods, in many ways these systems remain conceptually close to manually curated sets of rules (feature extractors). This might explain why LeadMine [8] (another contender), a purely rule-based system, outperforms most of the submitted ML-based counterparts. At the same time, it is possible to reduce manual interventions to the bare minimum by treating tokenisation, word encoding and feature extraction as subtasks in a global machine learning task, and this is exactly the kind of problems that deep artificial neural networks (ANNs) excel at. As we have already mentioned, neural networks can automatically learn morphology unaware word representations, and the same is true about morphology aware encodings. Furthermore, deep convolutional neural networks can automatically optimise feature extraction during training [14]. Most importantly, the labelling itself can be done by recurrent neural networks. Recurrent networks are naturally sequential and Turing-complete, extremely powerful in sequence-to-sequence (also known as seq2seq or many-to-many) modelling (including labelling) [15]. In an unreviewed paper by Rei et al. [16] the authors have experimented with deep-learning applications in NER on several datasets, including CHEMDNER. Some of their models used a bidirectional RNN for character-level word embedding combined with a variation of the attention technique used to choose between word-level and character-level embeddings, though the labelling itself was done by a CRF. Convolutional networks have also been used for biomedical NER. Zhu et al. [17] have applied a deep CNN to automatically infer local context-sensitive features fed into a CRF classifier. In an unreviewed article Chiu and colleagues have showcased a complete ANN-only design, based on a combination of convolutional and recurrent layers [18]. The model uses word-level and character-level token embeddings. While the former were pre-trained, the latter were optimised during training by transforming a word’s matrix of per-character linear embeddings into a single vector using a bidirectional RNN. Concatenated word-level and character-level embeddings were then fed into a CNN to extract local features. In contrast with the former examples, this model opted for a deep RNN instead of a CRF for sequence labelling. The authors claim state-ofthe-art performance on the datasets they’ve used, though quite unfortunately they have not tried to apply their model to a chemical dataset. All these examples make it self-evident that a pure ANN specifically targeting the CHEMDNER CEM subtask can perform as well (if not better) that conventional models, whilst relying on no imposed rules or databases whatsoever. Having set this as the main purpose of this study, we have developed a highly modular deep-learning model incorporating multiple novel features, including trainable targeted tokenisation.

## Materials and methods

### Problem formulation

We consider named entity recognition as a combination of two problems: segmentation and sequence labelling. Given:

- an ordered set of *N* character sequences *X* = (*X*_1_,…, *X_N_*), where *X_i_* = (*^i^c*1,…, *^i^ c_n_*) is a character sequence;
- an ordered set of *N* annotations *Y* = (*Y*_1_,…, *Y_N_*), where *Y_i_* is a sequence *Y_i_* = (*^i^y_1_,…, ^i^ y_n_*) and *^i^y_j_* is a tuple of two boolean labels (*^i^s_j_*,*^i^ e_j_*) showing whether the corresponding character is the beginning of a chemical entity and/or part of one, respectively;
 our task is to create a *predictor P: X → Ŷ*, where *Ŷ* is a set of inferred annotations similar to *Y*. We also introduce a *tokeniser* 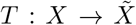, where 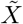 is an ordered sequence of character subsequences (tokens), thus slightly redefining the objective function to target per-token annotations. Provided that the *tokeniser* is fine enough to avoid tokens with overlapping annotations, this redefined problem is equivalent to the original one.

### Datasets

We used the CHEMDNER corpus [1] to train and validate our models. The corpus contains ten thousand abstracts from eleven chemistry-related fields of science with over 84k manually annotated chemical entities (20k unique) of eight types:

- ABBREVIATION (15.55%)
- FAMILY (14.15%)
- FORMULA (14.26%)
- IDENTIFIER (2.16%)
- MULTIPLE (0.70%)
- SYSTEMATIC (22.69%)
- TRIVIAL (30.36%)
- NO CLASS (0.13%)

The MULTIPLE class represents phrases containing several entities of other classes separated by non-chemical words. The CHEMDNER corpus comprises three parts: training (3.5k abstracts), development (3.5k) and testing (3k) datasets. We joined the first two datasets, randomly shuffled the result and separated 10% for a validation dataset to monitor overfitting. The other part of the split was used for training. We only used the official test dataset to estimate performance upon training completion.

### Design choices

**Deep-learning models**. We have utilised three types of neural networks: one-dimensional (1D) convolutional neural networks (CNN), recurrent neural networks (RNN) and time-distributed dense (fully-connected) networks (TDD). In their essence, one-dimensional convolutional neural networks are trainable feature extractors applied along a sequence evolving in time. A deep CNN [14] trains to extract time-invariant hierarchies of features at each time-step while optimising an objective function. Since texts are sequential, that is a value *t_i_* at time-step *i* can be conditioned on the previous and/or the following time-steps, a time-invariant model alone is not sufficient. We used recurrent neural networks – highly powerful trainable state machines theoretically capable of modelling relationships of arbitrary depth – to process CNN-extracted features. These networks train by back-propagating the error through time, which in deep sequences may lead to vanishing or exploding gradients. Several types of RNNs have been developed to better handle long-term dependencies, most notably the long short time memory network (LSTM) and gated recurrent unit network (GRU) [19]. Both architectures use trainable gates controlling the data flow and memory updates. The GRU architecture is a newer and lighter alternative to the widely adopted LSTM, with two trainable gates instead of the latter’s three resulting in less parameters to optimise, a desirable trait when training data are scarce. Comparative studies haven’t found any consistent performance advantages of either GRUs or LSTMs, though the former tend to converge faster [20]. To further improve performance, it is common to use bidirectional RNNs (biRNNs) “reading” sequences in both directions. Finally, we used a time-distributed fully connected network [21] (also known as dense networks or multilayer perceptrons) with the sigmoid activation function to generate label probabilities. In contrast with traditional bulky dense networks that process the entire input at once, TDDs apply a lightweight multi-layer perceptron (MLP) to each time-step in a sequence, drastically reducing the number of parameters and making it possible to analyse variable-length inputs.

**Stateful learning**. Texts come in all sizes, which is quite problematic for most machine-learning methods. Although one of the RNNs’ key selling points is their ability to naturally handle variable-size inputs, it’s hard to implement an RNN in a way that takes full advantage of this feature whilst staying computationally efficient. Two mainstream solutions exist. The most natural – and arguably the least computationally efficient – solution implies grouping and encoding (i.e. representing as numeric tensors) equally sized texts together. This method introduces a lot of extremely small sample batches greatly increasing gradient variance and, by extension, hindering model convergence. Alternatively, one can use zero-padding (artificially increasing length by appending zeros to numerically encoded sequences). This procedure greatly increases sparsity and the memory overhead, because full-sized texts can vary greatly in length. It is thus more efficient (and common) to break texts into individual sentences. Despite being more computationally efficient, this method is less flexible, because it introduces a sentence length limit and requires a sentence segmentation model. It also strips aways text-wide context. Quite fortunately, there is another relatively novel technique known as stateful learning. Although it has not yet gained any noticeable adoption in the community (partly due to complicated data handling described below), it combines the best of both aforementioned methods: no text-size restrictions, no sentence segmentation model dependencies and negligible memory overhead. Normally RNNs only keep their state within a single batch of samples and reset it between batches, because there is no guarantee that the next batch is somehow related to the previous one. In contrast with conventional setups, an RNN configured for stateful learning treats a sample (row) *j* in batch *i* + 1 as the direct continuation of sample *j* in batch *i*, making it possible to break long sequences into fixed-size windows without resetting the context when moving from one batch to another. Simply put, stateful learning allows RNNs to transcend the batch barrier and, in theory, keep track of the context as long as required. To make it technically possible, the batch-size must be fixed at construction time and the data must be preprocessed to satisfy the aforementioned property. We ve developed a bin packing-based data preprocessing algorithm to achieve this goal. Given a batch size of *n*, we distribute input texts into *n* bins while trying to keep the accumulated lengths equal between all bins. We then concatenate texts inside each bin into super-sequences, stack them and break into chunks of *l* time-steps. This procedure is easily reversible, making it possible to recover annotations for individual texts. Additionally, since in bidirectional RNNs it only makes sense to keep track of the forward-evolving state, we have developed a “half-stateful” bidirectional RNN wrapper layer (HS-biRNN) that takes care of forward inter-batch state transfers and can be used with any RNN architecture (e.g. LSTM or GRU).

### Text preprocessing

We have done no text-preprocessing except for tokenisation. Accurate tokenisation is highly important in token-level NLP tasks [5]. On the one hand, this process can isolate semantically and morphologically stable character sequences, making it easier for the model to focus on the data. On the other, tokenisation may lead to overlapping annotations if the rules fail to separate several adjoint entities or non-entity characters from entities. Most popular tokenisers rely on a hierarchy of rules optimised for standard English, though there are some specifically designed for biomedical and chemical texts. For example, the tokeniser implemented in the ChemDataExtractor package [22] overrides some rules in the Penn Treebank policy to better handle chemical entities:

> Tokens are split on all whitespace and most punctuation characters, with exceptions for brackets, colons, and other symbols in certain situations to preserve entities such as chemical names as a single token.

In other words, as with any rule-based technique, it’s notoriously hard to create an optimal tokeniser equally adequate for recovering standard vocabulary and diverse chemical entities, because they have different underlying morphology – a tokeniser has to be context-aware. We believe that instead of trying to manually create a general-purpose tokeniser one can use an alternative specifically trained to accurately recover target entities. Such a tokeniser will only be used to preprocess text for a subsequent NER model alleviating the need to recover irrelevant words. Since we have found little to no research on trainable tokenisers, we have developed our own model based on a “break and stitch” strategy: a primary extra-fine segmentation followed by a refinement step trying to recover target entities (Fig.1). We have used the following Perl-style regular expression to carry out the first step: \w+|[^\s\w]. The expression groups together Unicode word characters (i.e. most characters that can be seen in a word in any language, including numbers) and separates all other characters. For example, it breaks **2-amino-1-methyl-6-phenylimidazo[4,5-b]pyridine** into nineteen fragments: **2, –, amino, –, 1, –, methyl, –, 6, –, phenylimidazo**, [, **4,,, 5, –, b**,], **pyridine**. As expected, the result is heavily over-fragmented. On the bright side, our analyses of the tokenised CHEMDNER corpus showed a near-complete absence of tokens overlapping several entities, making it possible to accurately reconstruct large entity tokens by stitching several fragments together. To detect stitch points we have designed a lightweight stateful sequence-to-sequence CNN-RNN model processing raw untokenised text. The model consists of a linear character encoding layer, two consecutive 1D CNN layers, each with 256 (3-characters wide) filters, followed by two half-stateful bidirectional GRU layers (32 cells each) and a time-distributed sigmoid classifier that outputs a binary tag for each character in the sequence. Positive tags mark stitch points. We have used the same training and validation splits to train this tokeniser alongside the NER model.

**Figure 1.**
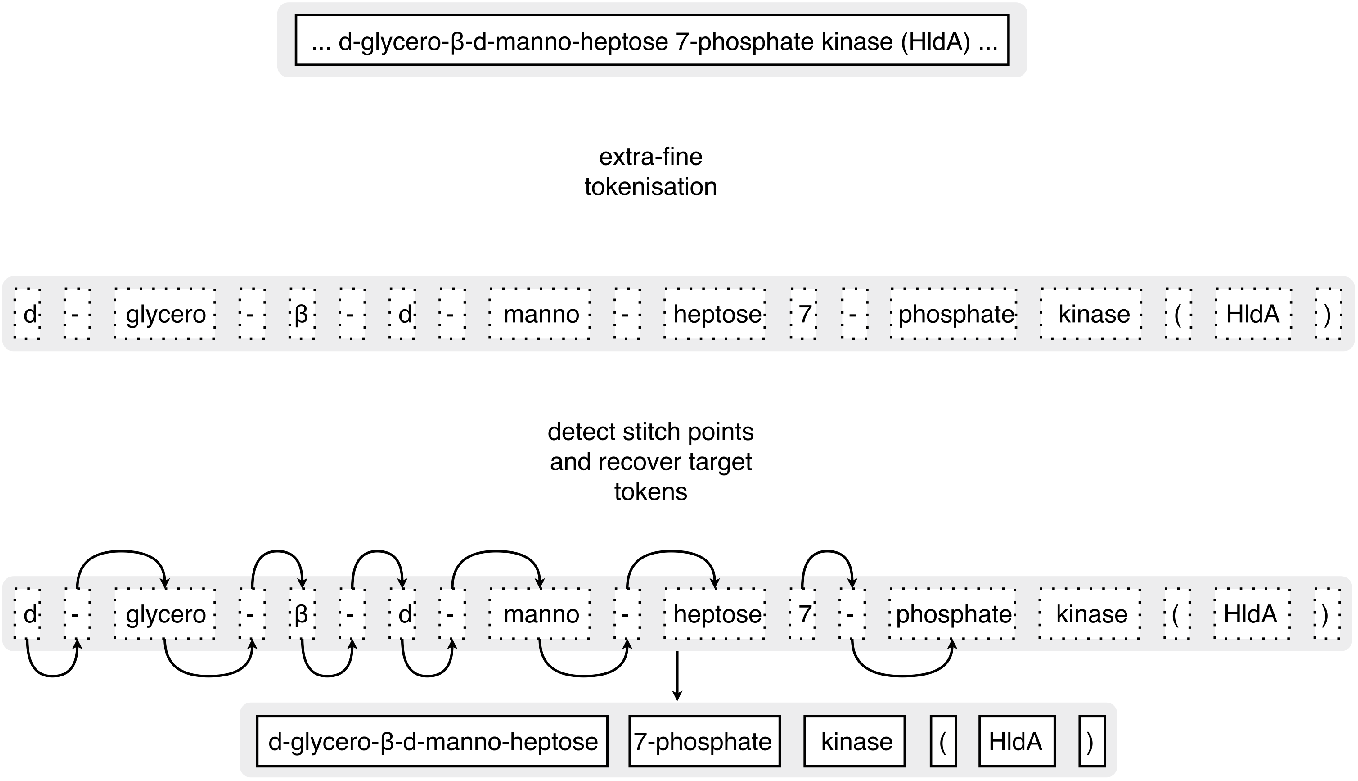
Text tokenisation. The break and stich targeted tokenisation strategy employed by our trainable tokeniser.

### The NER model

All the models we trained had two input nodes: one for pretrained word-level embeddings and another one for encoded token strings. The strings were encoded as integer vectors containing character identifiers. We trained 300-dimensional Glove embeddings with default configurations [23] on a corpus of 5 *·* 10^5^ random PubMed abstracts from the same categories as the CHEMDNER abstracts: BIOCHEMISTRY & MOLECULAR BIOLOGY, APPLIED CHEMISTRY, MEDICINAL CHEMISTRY, MULTIDISCIPLINARY CHEMISTRY, ORGANIC CHEMISTRY, PHYSICAL CHEMISTRY, ENDOCRINOLOGY & METABOLISM, CHEMICAL ENGINEERING, POLYMER SCIENCE, PHARMACOLOGY & PHARMACY and TOXICOLOGY [1]. Character-level embeddings were optimised during training using the same approach described in [18]. This block consisted of a trainable linear character-embedding layer transforming vectors of character codes into matrices of 32-dimensional character embeddings. These word matrices are then processed by a standard biGRU (16 cells) layer producing a 32-dimensional vector per token [24].

Instead of concatenating word- and character-level embeddings before feeding them into a single CNN or RNN block, we used separate two-layers deep 1D CNNs for each embedding type to increase the number of degrees of freedom without using too many convolutional filters. Features extracted by these independent blocks were subsequently concatenated and fed into a two-layers deep HS-biGRU. The network then bifurcates again:

1. The first branch continues with an additional HS-biRNN followed by a time-distributed sigmoid layer. The layer outputs the probability that a given time-step is a part of a named entity.
2. The second one starts with an arithmetic node multiplying the probabilities produced by the other branch and the output from the preceding HS-biGRU block at each time-step. The result is then fed into a single HS-biRNN layer followed by a time-distributed sigmoid layer, yielding the probability that a given time-step is the beginning of an entity.

First of all, it’s important to show that this labelling method remotely resembles the widely used IOB scheme with three mutually exclusive labels: Inside/Outside/Beginning (of an entity) [25]. At the same time, in contrast to this scheme (or any other scheme with mutually-exclusive tags), our labels are not mutually exclusive and are codependent at the same time due to the multiplication node. Since we try to minimise both predictors their error is back-propagated through the graph, creating a reinforcing loop with two effects: (1) it theoretically encourages the part-detected to better pay more attention to single-token entities and (2) it helps the beginning-detector attend to entity parts.

Here we have described all design elements of the final (fully-featured) NER model. We also examined the impact of several large-scale changes on its performance. In particular, we have compared GRU and LSTM architectures and tried replacing the CNNs with additional recurrent layers. More importantly, we have compared stateful and conventional biRNNs trained unsegmented full-sized texts and stacked sentences respectively. We used the GeniaSS [26] sentence segmentation model to carry out this comparison. Having limited computational resources and time constraints we have not tried to fine-tune any hyper-parameters: all convolutional layers comprised 256 (3 time-steps wide) filers, and all HS-biRNN layers contained 32 recurrent cells.

### Training and testing

The project was implemented in Python 3.5 using deep-learning frameworks Keras 2[27] and TensorFlow 1.3 [28]. All computations have been carried out on an Ubuntu Linux server with two Intel Xeon E5 CPUs (10 cores and 20 threads each), 512GB of RAM and four Nvidia Titan X GPUs. We used the Adam optimiser [29] with default parameters recommended by the authors. The networks were trained for 40 epochs with a callback saving weights upon improvements in performance on the validation dataset.

During testing, we specifically targeted the CHEMDNER chemical entity mention (CEM) subtask. Since deep-learning models are inherently non-deterministic due to random weight initialisation and stochastic optimisation, we have evaluated each design variation by averaging estimated probabilities from 10 independently trained networks (as in [16]). To add some perspective, we report all models that have achieved a CEM F-score of 80% and above in the CHEMDNER challenge1, though their current accessibility is worth mentioning. The models introduced by teams 184, 185, 192 either have not been published at all or the links have become inactive. LeadMine (179) [8] is exclusively commercial. While there is a GitHub repository for the model devised by Lu et al. (team 231) [9, 30], it literally contains nothing but a link to an archive (which supposedly contains the model), uploaded to a file-hosting service that requires a proprietary application to download the archive. Since both the file-hosting and the application are only available in Chinese, we have been unable to download the archive and thus consider it inaccessible. Becas (team 197)[31] and tmChem (team 173) [2] both provide different web-based APIs, making it possible to submit texts to annotation servers or, in case of tmChem, even download precomputed annotations for PubMed abstracts. With tmChem there is also an option to build the tool from sources, though the source archive does not seem to come with a trained model, because our stand-alone installation has produced random annotations. Both Chemspot (team 198) [32] and BANNER-CHEMDNER (team 233) [33] are available for stand-alone installation from sources.

**Table 1.** Performance scores for the CHEMDNER chemical entity mention (CEM) subtask. CHEMDNER challenge team IDs are given in parenthesis in the Model column (where available; performance scores for these models have been taken from Table 4 in [30]). We provide ChemDataExtractor performance scores reported by the authors.

| Model | Precision % | Recall % | F1-score % |
| --- | --- | --- | --- |
| Our model | 88.6 | 88.8 | 88.7 |
| ChemDataExtractor [22] | 89.1 | 86.6 | 87.8 |
| tmChem (173) [2] | 89.2 | 85.8 | 87.4 |
| (231) [9] | 89.1 | 85.2 | 87.1 |
| LeadMine (179) [8] | 88.7 | 85.1 | 86.9 |
| (184) | 92.7 | 81.2 | 86.6 |
| Chemspot (198) [32] | 91.2 | 82.3 | 86.7 |
| Becas (197) [31] | 86.5 | 85.7 | 86.1 |
| (192) | 89.4 | 81.1 | 85.1 |
| BANNER-CHEMDNER (233) [33] | 88.7 | 81.2 | 84.8 |
| (185) | 84.5 | 80.1 | 82.2 |

Apart from the models submitted for the CHEMDNER challenge we have also considered ChemDataExtractor [22], a recently introduced general purpose package for chemical text analyses, because its NER model is very much akin to [9], which is unavailable. Both utilise unsupervised word-clustering and CRFs, though ChemDataExtractor uses a hierarchical detection system with a built-in database updated in an online fashion to help it extract abbreviations and identifiers. ChemDataExtractor comes with a highly user-friendly Python API making it extremely easy to install and utilise. More importantly its overall combined CEM F-score of87.8% puts it on top of all models submitted for the CHEMDNER challenge.

## Results and discussion

Tokenisation, overlapping annotations and sequence lengths

We have processed the entire CHEMNDER testing dataset and searched for entities with overlapping annotations. Out of 25347 annotated entities in the testing dataset less than 0.19% spanned the same token, which is truly negligible. At the same time the tokeniser had a recall of 91.75% and precision of 93.32%. Therefore, it is able to accurately recover most of the annotated entities.

## Performance

First of all, in terms of time required to complete one training epoch the reference network (fig.2) incorporating stateful biRNNs trained over two times faster than its sentence-based sibling with conventional biRNNs and had a lighter memory footprint. We observed no significant impact on the F-score, though. There was no observable advantage in using LSTM cells over GRUs, either. On the contrary, GRUs trained and converged faster and showed slightly better performance on the testing dataset. Convolutional layers were crucial for good performance. On average, replacing the CNN-layers with one or two hs-biGRU layers reduced the F-score by ∽ 1.5 − 2.3% and hampered the training process.

**Figure 2.**
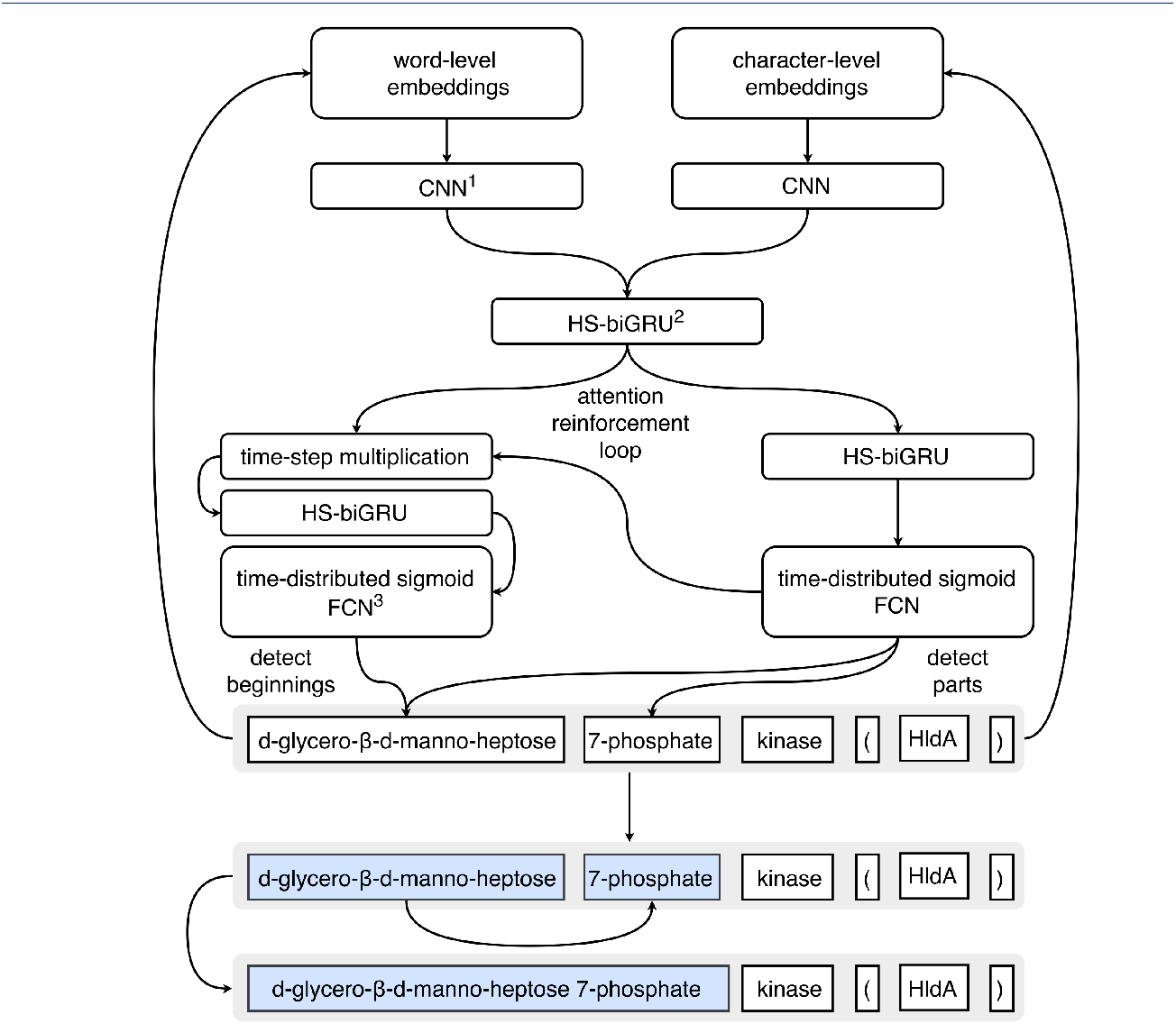
Model architecture. The figure illustrates the topology and of the best-performing full-featured model. ^1^CNN – convolutional neural network ^2^HS – biGRU – half-stateful bidirectional gated recurrent unit ^3^FCN – fully-connected network

On the CHEMDNER CEM subtask our fully-featured network has gained the F-score of 88.7%. Therefore, it outperforms all models submitted for the CHEMDNER task by a significant margin, though the edge over ChemDataExtractor is less impressive (see table 1 for more details). Considering the inter-annotator agreement score of 91%, the model demonstrates near-human performance. Since the model does not discriminate between entity types, there is no way to calculate per-class precision values and, by extension, F-scores. Nevertheless, following Krallinger et al.[30] we report per-class precision in table 2. It’s important to note that following the CHEMDNER CEM evaluation rules we have only considered perfect matches. It is immediately clear that the model struggles greatly with rare entity types, i.e. NO CLASS and MULTIPLE, and excels at systematic and trivial names. Considering how rare the MULTIPLE type entities are (195 entities) and that they span several standard English words intertwined with different chemical entity types, this subpar performance is not surprising and is actually consistent with that of other tools reported in [30] (Additional file 3).

**Table 2.**
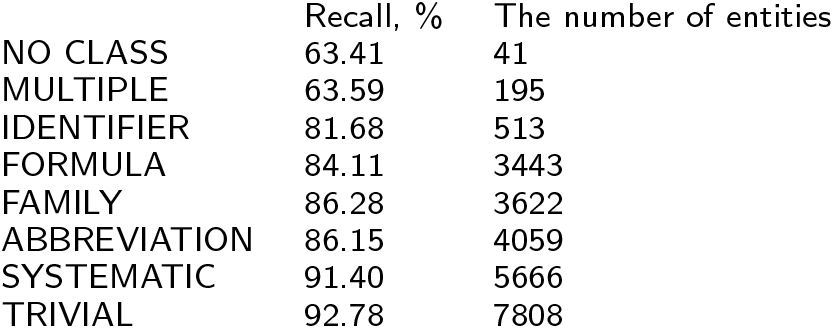
Recall values estimated for individual entity types. Only perfect matches were considered correct.

## Accessibility and the user interface

While analysing the NER systems submitted for the CHEMDNER task, we have found that neither the source code, nor the trained models are available for some of the best-performing tools, limiting the ability to use and validate them. We thus made it our priority to publish all the source code needed to train and use our models. All materials are openly available on GitHub [34] in a separate frozen branch (chemdner-pub) of a natural language processing package SciLK. While the package itself, being in the early stages of development, is bound to change, the separate branch will retain the version required for these models to work. In addition to the core library, the branch contains Jupyter notebooks with code and commentaries sufficient to reproduce our work, i.e. train a tokeniser and a named entity detector, a notebook with usage demonstration and an archive with trained models. A standard dual-core laptop with 8GB of RAM should be sufficient to use the models for inference. While the software should theoretically work under Microsoft Windows, we have only tested it on machines running Mac OS X and GNU-Linux. Should a user want to train a similar model, we recommend doing it on a machine with at least 32GB of RAM and a graphics processing unit (GPU) with at least 8GB of VRAM. Although a GPU is not strictly required for training, it takes roughly twice as much time to train a fully-featured model on a 20-core CPU-only system.

## Conclusions

Here we have presented our deep-learning model for chemical named entities recognition in biomedical texts, trained and evaluated on the CHEMDNER corpus. Given its high performance, the model proves that chemical named entity recognition can be done efficiently with no manually created rules or curated databases whatsoever. We also showcase several novel or rarely used approaches and design choices that, to the best of our knowledge, have never been used in biomedical or chemical NER. Most notably, we advocate the use of specialised trainable tokenisers and stateful recurrent neural networks. Nevertheless, we clearly see several directions for further improvement. For one, due to time constraints we have not investigated many hyper-parameter and topology options. Secondly, while avoiding complicated preprocessing has been one of the top-priorities, we still believe that additional information that cannot be extracted from the CHEMDNER corpus itself can further increase performance. In particular, we think that part of speech tags or other external annotations can greatly benefit the system. We also think that much more research should be done on targeted tokenisers, considering that our tokeniser had a rather primitive design.

## Author’s contributions

Ilia Korvigo and Mikhail Skoblov conceived and curated the study. Ilia Korvigo, Maxim Holmatov and Anatolii Zaikovskii contributed to data acquisition and supervision. Model development, training and validation was performed by Ilia Korvigo. Software development was performed by Ilia Korvigo. The manuscript was written by Ilia Korvigo and Maxim Holmatov.

## Acknowledgements

This work was carried out in collaboration between the Laboratory of Functional Analysis of the Genome (Moscow State Institute of Physics and Technology, Moscow, Russia) and the Laboratory of Microbiological Monitoring and Bioremediation of Soils (All-Russia Institute of Agricultural Microbiology, St. Petersburg, Russia).

